# Mutational impact of culturing human pluripotent and adult stem cells

**DOI:** 10.1101/430165

**Authors:** Ewart Kuijk, Myrthe Jager, Bastiaan van der Roest, Mauro Locati, Arne Van Hoeck, Jerome Korzelius, Roel Janssen, Nicolle Besselink, Sander Boymans, Ruben van Boxtel, Edwin Cuppen

## Abstract

Genetic changes acquired during *in vitro* culture pose a potential risk for the successful application of stem cells in regenerative medicine. To assess mutation accumulation risks induced by culturing, we determined genetic aberrations in individual human induced pluripotent stem cells (iPS cells) and adult stem cells (ASCs) by whole genome sequencing analyses. Individual iPS cells, intestinal ASCs and liver ASCs accumulated 3.5±0.5, 7.2±1.0 and 8.4±3.6 base substitutions per population doubling, respectively. The annual *in vitro* mutation accumulation rate of ASCs adds up to ∼1600 base pair substitutions, which is ∼40-fold higher than the *in vivo* rate of ∼40 base pair substitutions per year. Mutational analysis revealed a distinct *in vitro* induced mutational signature that is irrespective of stem cell type and distinct from the *in vivo* mutational signature. This *in vitro* signature is characterized by C to A changes that have previously been linked to oxidative stress conditions. Additionally, we observed stem cell-specific mutational signatures and differences in transcriptional strand bias, indicating differential activity of DNA repair mechanisms between stem cell types in culture. We demonstrate that the empirically defined mutation rates, spectra, and genomic distribution enable risk assessment by modelling the accumulation of specific oncogenic mutations during typical *in vitro* expansion, manipulation or screening experiments using human stem cells. Taken together, we have here for the first time accurately quantified and characterized *in vitro* mutation accumulation in human iPS cells and ASCs in a direct comparison. These results provide insights for further optimization of culture conditions for safe *in vivo* utilization of these cell types for regenerative purposes.

## Introduction

As infinite supplies of undifferentiated and specialized cells, induced pluripotent stem (iPS) cells and adult stem cell (ASC)-derived organoids hold great potential in regenerative medicine^1^. Furthermore, stem cells have become invaluable tools in pharmacology and toxicology for *in vitro* testing. Maintenance of genomic integrity in culture is a prerequisite for the successful application of stem cells, as unwanted mutations may influence drug responses and toxicology measurements or might lead to oncogenic transformation following transplantation. However, multiple studies have identified recurrent genomic alterations that originate in routinely cultured human adult and pluripotent stem cell lines^2–15^. Although these studies suggest that the genomic integrity of stem cells is reduced in culture, the majority of these studies have been performed on bulk cultures with methods that have limited resolution. Analysis of bulk samples poses important limitations as only mutations that are shared among the majority of the cells are detectable. This results in a bias towards those genomic alterations that confer a selective advantage upon the cells or those that occurred early in the culturing process. The mutational impact of the culturing process itself and the responsible mutational mechanisms remain as yet unclear. A better understanding of the mutational processes that are active in culture might enable us to improve genomic stability in culture, e.g. by changing the culture conditions.

We recently established a method to accurately identify *in vivo* acquired somatic mutations in individual stem cells at base pair resolution by combining whole genome sequencing (WGS) with *in vitro* clonal expansion. In short, stem cells are seeded as single cells and propagated to obtain sufficient cells for DNA isolation and subsequent whole genome sequencing (WGS). Bioinformatic analyses are performed that identify with high confidence those mutations (single nucleotide variants, indels, copy number alterations (CNAs), structural variants (SVs), aneuploidies) present in the original cell and filter out subclonal mutations that occurred after the single cell step ^16^. With this approach, high noise rates of *in vitro* amplification approaches are avoided and mutation loads as low as hundreds of mutations genome-wide can be reliably detected. The resultant genome-wide mutation spectra and distribution were shown to provide novel insights into the activity of specific mutational and DNA repair processes in adult stem cells *in vivo* and CRISPR/Cas9-edited organoids ^17–19^.

Here we have adapted this approach to systematically measure the mutational impact of *in vitro* culture on individual cells of three human stem cell types that were chosen for their relevance in pharmacology, toxicology and regenerative medicine: iPS cells, liver ASCs, and intestinal ASCs. Our study demonstrates that *in vitro* culture results in increased mutation rates and leaves a distinct but common mutational footprint related to oxidative stress in all three stem cell types, in addition to stem cell-specific mutational signatures. Furthermore, we used the measured quantitative and qualitative mutational characteristics to model genome wide mutation accumulation and perform genetic risk assessments associated with *in vitro* and *in vivo* applications of stem cells.

## Results

### Mutation accumulation in iPS cells and ASCs during culture

To investigate the mutational consequences of standard culture conditions on the genome of stem cells in an unbiased manner, we first established clonal human iPS and ASC lines. Each clonal line was cultured for ∼2–5 months (supplementary table 1), in which mutations were allowed to accumulate. Subsequently, a second clonal step was performed to determine the acquired mutations at the individual cell level. Clonal cultures and a matched nonclonal reference sample were subjected to WGS (supplemental figure 1). All germline variants and variants that accumulated before the first clonal step were excluded based on the reference sample. Variants that arose *in vitro* after the second clonal step will be subclonal and could be filtered out bioinformatically based on low variant allele frequency (VAF). This approach allowed us to specifically measure the mutations that accumulated between the two clonal steps. *In vitro* mutation accumulation was determined in 3, 4, and 6 subclones for the iPS cells, liver ASCs and intestinal ASCs, respectively.

The total number of stem cells that is required for a particular application like e.g. cell therapy or drug study is achieved after a certain amount of population doublings, which is defined by the cell division rate and the cell death rate. Thus, from a practical perspective, knowledge on the mutation rates per population doubling is more informative than the number of mutations per time unit. Therefore, we determined the population doubling rate for all cell types under the same conditions that were also used for the mutation accumulation experiments. We found a population doubling time of ∼23h for human iPS cells which is approximately twice as fast as for human liver and intestinal ASCs, that have population doubling times of ∼46h and ∼44h, respectively (figure 1a).

**Figure 1:**
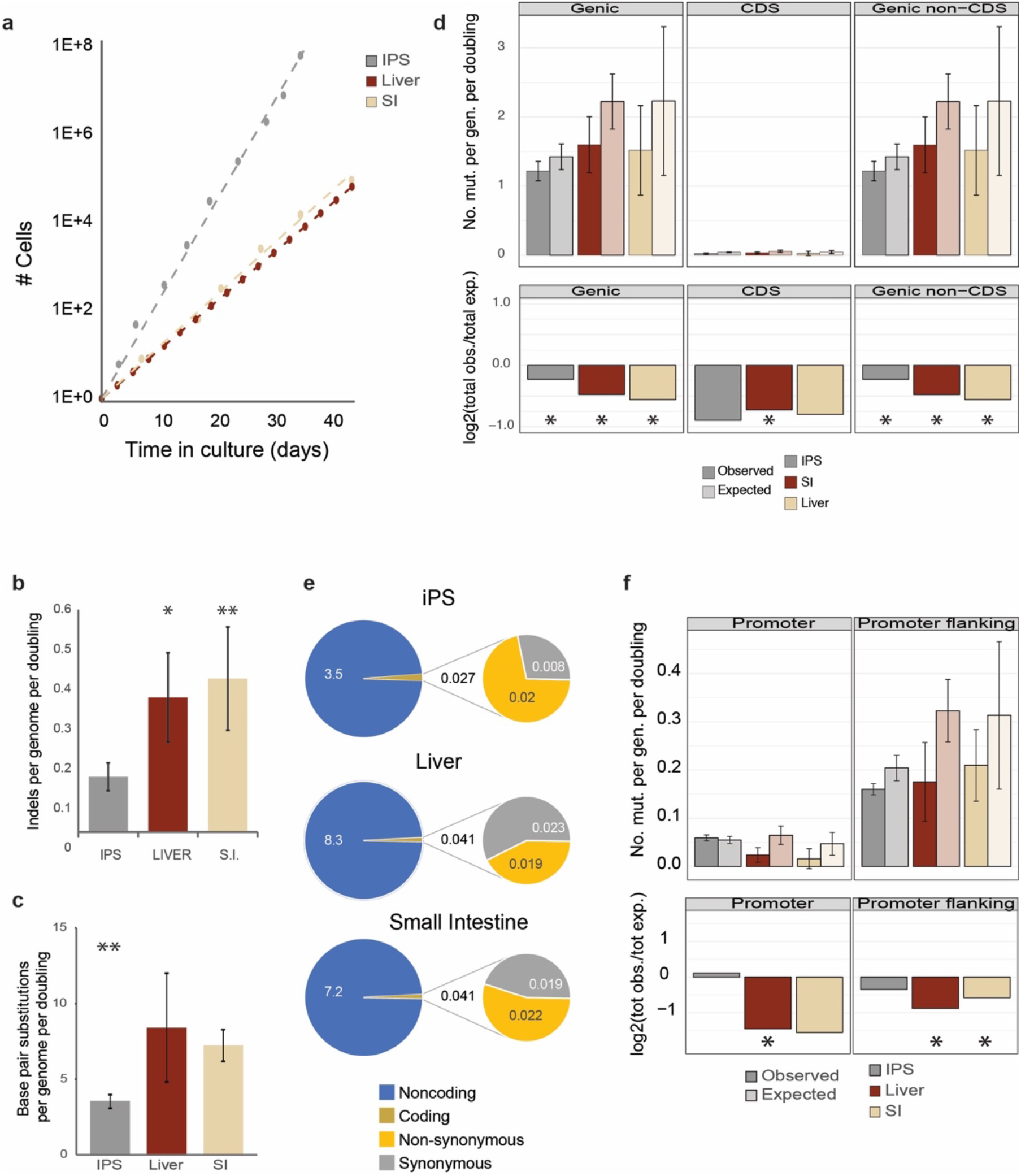
*In vitro* mutation accumulation in iPS cells, liver stem cells, and intestinal stem cells. **a**, Growth curves for iPS cells, liver stem cells, and intestinal stem cells **b**, Number of indels per genome per population doubling, asterisks denote statistically significant differences (ANOVA) **c**, Number of mutations per genome per population doubling, asterisks denote statistically significant differences (ANOVA) **d**, **T** op panels: genomic distribution of the observed versus the expected amount of mutations per genome per doubling at genic regions, coding sequences and non-coding sequences. Error bars represent the standard deviation. Bottom panels: log2 of the ratio between the total number of observed mutations versus the total number of the expected amount of mutations at genic regions, coding sequences and non-coding sequences. Asterisks denote statistically significant differences between observed and expected (binomial test). **e**, Total number of somatic base substitutions per cell type (left circles) and those affecting protein-coding DNA (right circles), **f**, Top panels: genomic distribution of the observed versus the expected amount of mutations per genome per doubling at promoter and promoter flanking regions. Error bars represent the standard deviation. Bottom panels: log2 of the ratio between the total number of observed versus the total number of the expected amount of mutations at promoter and promoter flanking regions. Asterisks denote statistically significant differences between observed and expected.

The karyotypes of all clones and subclones for all three stem cell types were analyzed by sequencing read-depth coverage and found to be normal without any gross chromosomal abnormalities. CNAs were not observed for any of the subclones. Furthermore, no structural variants (SVs) were observed for the iPS cells. In the ASCs, however, we did observe seven deletions and one inversion (supplementary figure 2 and supplementary table 2). In one of the liver ASC subclones, a 139 bp deletion was detected and six deletions (ranging from 51 to 97,064 bp) were observed in four of the intestinal stem cell subclones. Additionally, one of the intestinal subclones also contained an inversion of 4,084 bp. Four of the deletions involved known common fragile sites. In total, four genes overlapped with the SVs, but the impact on gene function is considered low as none of the SVs involved coding sequences (supplementary table 2).

In total, we detected 201 small insertions and deletions (indels) in the 13 subclones (supplementary table 1). After correction for the stem cell-specific growth rates, iPS cells, acquired 0.15 indels per genome per population doubling, which is statistically significantly different from the liver and intestinal ASCs with 0.36 and 0.41 indels per genome per population doubling respectively (ANOVA, p = 0.03; figure 1b). To assess the functional impact on proteins, we performed PROVEAN^20^ analysis, which revealed that three indels are in protein coding sequences, of which one in an intestinal ASC clone is predicted to be deleterious for protein function (supplementary table 3).

We identified a total of 3,888 base pair substitutions unique to the 13 subclones. Liver ASCs acquired 8.4±3.6 base pair substitutions per genome per population doubling. In intestinal ASCs, the mutation accumulation rate was similar with 7.2±1.0 base pair substitutions per genome per population doubling. The number of base pair substitutions was lower in the iPS cells than in the ASCs, with 3.5±0.5 mutations per genome per population doubling (ANOVA, p = 1.5E^-04^; figure 1c).

At the observed *in vitro* mutation rates, individual iPS cells accumulate 1,328±171 mutations per year, intestinal ASCs accumulate 1,426±207 mutations per year, and liver stem cells accumulate 1,608±689 mutations per year. For the ASCs, these values are ∼40-fold higher than the *in vivo* mutation accumulation rate of ∼40 base pair substitutions per year ^18^.

Next, we analyzed the potential functional consequences of the observed mutations. Base pair substitutions were significantly depleted in genic regions for all three stem cell types (figure 1d). The depletion was not restricted to the protein coding sequence, but also included non-coding sequences, indicating that the depletion in genic regions is mainly caused by enhanced repair activity in genic regions and not by selection against deleterious mutations. The depletion was less pronounced for the iPS cells with 0.027 mutations in coding sequences per genome per population doubling compared to 0.041 in both ASC types (figure 1e). In total, we identified 14 non-synonymous mutations across all samples, 5 in the iPS cells, 2 in the liver ASCs, and 7 in the intestinal ASCs. For each stem cell type, this was found to convert in ∼0.02 non-synonymous mutations per genome per population doubling (figure 1e and supplementary table 4). None of the non-synonymous mutations affected known cancer genes based on the census of human cancer genes^21^ or have previously been described to confer a selective advantage over stem cells in culture ^6,14^. We are therefore confident that the observed mutations provide an unbiased insight into mutation accumulation free of selection.

Mutations in promoter regions can affect gene activity and thereby contribute to disease ^22,23^. Mutations were depleted in the promoter regions^24^ of both ASC types (figure 1f). No depletion was observed in the promoter regions of the iPS cells. Mutations in heterochromatin are generally considered harmless, because the risk that function elements are affected is low. In all three stem cell types, mutations were significantly enriched in heterochromatic laminin associated domains (LADs)(supplementary figure 3), as is the case for mutations that arise during reprogramming ^25^. Functionally relevant genomic regions thus appear more protected from mutation accumulation in all three stem cell types, while heterochromatic LADs are more susceptible to mutation accumulation.

### Mutational patterns of the SNVs

To obtain insight into the mutational processes induced by culturing, we analyzed the mutation spectra in more detail. In line with our previous observation on liver stem cells^26^, C to A transversions were the predominant base substitutions in the mutational spectrum of all three stem cell types, encompassing nearly 30% of the base substitutions in the liver ASCs and over 35% of all the base substitutions in the iPS cells and the intestinal ASCs (figure 2a). This mutation type has been linked to reactive oxygen species (ROS)^27^. In contrast with mutation accumulation in intestinal stem cells *in vivo ^18^*, we observed only a limited relative contribution of C to T changes in a CpG context in intestinal stem cells in culture (figure 2a). The low contribution of this mutation type in *in vitro*-cultured intestinal stem cells indicates that other mutational processes are more dominant. Transcriptional strand-bias was observed for T to A and T to C changes in the intestinal ASCs and for T to C changes in the iPS cells (supplemental figure 4).

Next, we compared the mutational profiles of the *in vitro* cultured stem cells to the COSMIC signatures^17^. To confirm whether the mutational processes are indeed different *in vivo*, we also included the previously described mutational profiles of *in vivo* accumulated mutations for the liver, the small intestine and the colon in this analysis ^18^. In line with previous observations, hierarchical clustering based on similarity with the COSMIC signatures resulted in a clear segregation between the spectra of *in vivo* acquired mutational signatures versus *in vitro* acquired mutations (figure 2b). Strikingly, COSMIC signature 18 contributed to all three stem cell types (figure 2b and supplemental figure 5). Although the etiology of this signature is as yet unknown, signature 18 is characterized by a large contribution of C to A changes, in line with our observation that this is the dominant base substitution in all three stem cell types. Hence, *in vitro* mutation accumulation seems to be mainly determined by culture-induced high levels of oxidative stress resulting in largely similar mutational patterns irrespective of stem cell type.

**Figure 2:**
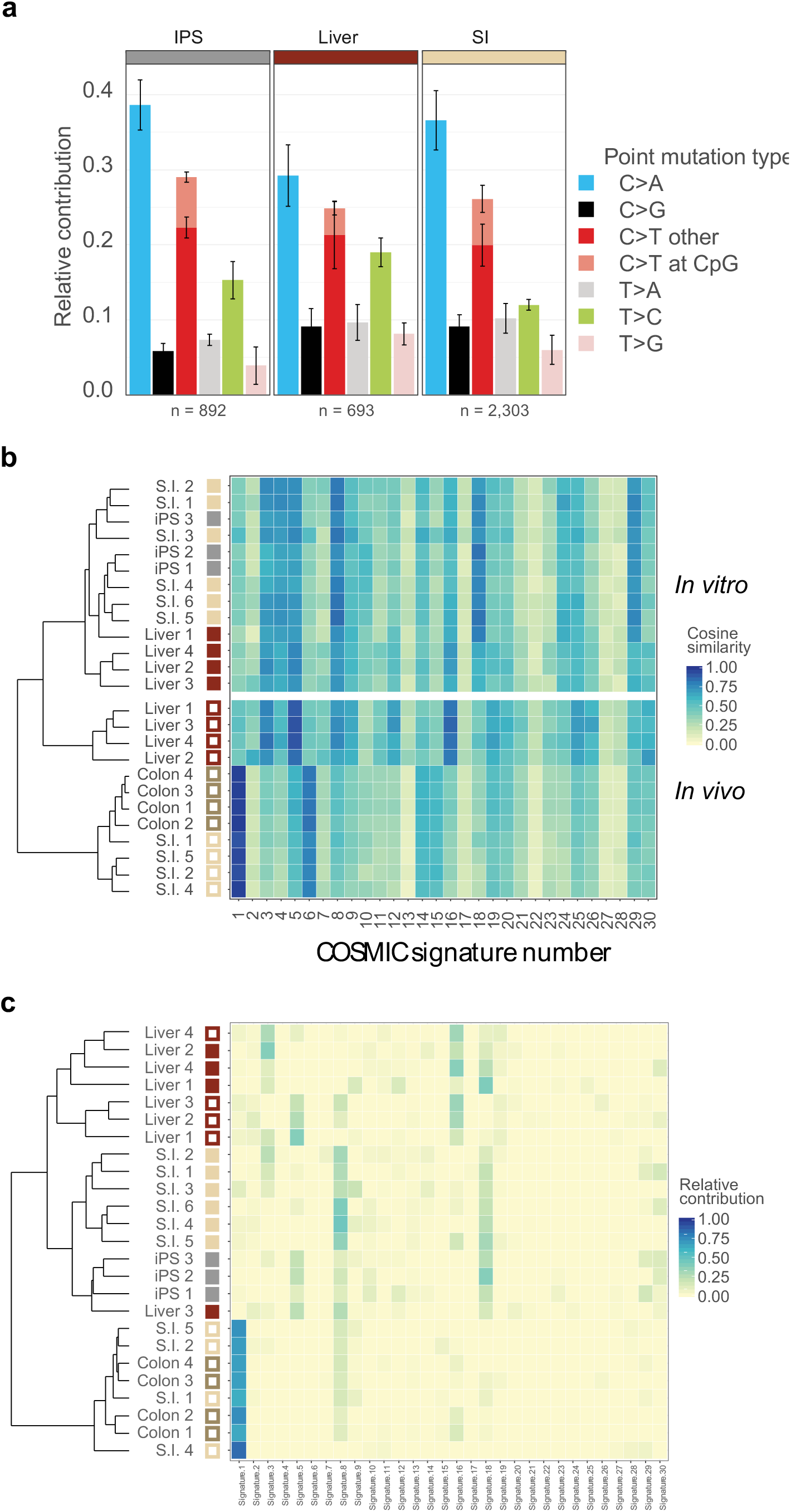
Mutational spectrum and signature analysis. **a**, Relative contribution of the indicated base substitution types to the mutation spectrum. Per stem cell type, data are represented as the mean relative contribution of each mutation type over all subclones. Error bars represent the standard deviation. The total number of SNVs is indicated. **b**, Cosine similarity heatmap between the mutational profiles in the subclones and the COSMIC signatures, using unsupervised clustering of the subclones. Open squares indicate *in vivo* clones from Blokzijl et al. ^18^, closed squares indicate *in vitro* subclones (this study). **c**, Relative contribution of COSMIC signatures to the mutational profiles of the subclones using unsupervised clustering of the subclones. Open squares indicate *in vivo* clones from Blokzijl et al. ^18^, closed squares indicate *in vitro* subclones (this study).

While COSMIC signatures 1 and 6 have a relatively large contribution to mutation accumulation in the intestine *in vivo*, their contribution to the *in vitro* mutation spectra is limited. Likewise, the contribution of signature 5 to mutation accumulation in liver stem cells is more prominent *in vivo* than *in vitro*. Together, these findings demonstrate that *in vivo* mutational processes only play minor roles in *in vitro* mutation accumulation.

Finally, we compared the relative contribution of all COSMIC signatures^17^ to the mutational signatures of each subclone, followed by hierarchical clustering based on these contributions. Cells largely clustered by stem cell type, in particular the iPS and intestinal stem cells, indicating that stem-cell specific mutational processes are still active in these *in vitro* cultured stem cells (figure 2c). Signature 5 contributed to the mutational profile of the iPS cells, but not the majority of the *in vitro*-cultured ASCs (figure 2c and supplemental figure 5). Signature 5 is characterized by transcriptional strand bias for T to C substitutions, as was observed for the iPS cells (supplemental figure 4). Signature 8 contributed to all small intestinal stem cell clones and, to a lesser extent, to the iPS cells (figure 2c and supplemental figure 5). Signature 8 did not contribute to 3 out of 4 liver stem cell clones.

### The mutational risk of *in vitro* culture

The *in vitro* mutational processes may lead to the transplantation of cells carrying pathogenic mutations. We therefore used the empirically defined stem-cell specific mutation rates, mutation spectra, and genomic distribution to model the risk of oncogenic mutations to occur during *in vitro* culture. This revealed a near linear correlation between the cumulative number of pathogenic mutations in oncogenes ^28^ as a function of the number of stem cells that are generated *in vitro*(figure 3a). In liver ASCs there is one oncogenic mutation per 2.7*10^6^ cells, in intestinal ASCs one per 4.6*10^6^ cells and in iPS cells one per 6.8 *10^6^ cells. When 10^8^ intestinal ASCs are produced *in vitro* (a high-end estimate required for a cellular transplantation), the chance for at least one oncogenic mutation in the population is approximately 1. To place this chance in perspective we also calculated the number of oncogenic mutations that occur in the colon *in vivo*. The risk for 10^8^ *in vitro* produced intestinal ASCs is equivalent to the risk of accumulating an oncogenic mutation in any stem cell in human colon *in vivo* in 100 days (figure 3b). Because not all possible oncogenic mutations are relevant for all three stem cell types we also looked at more specific mutations. For example, in human pluripotent stem cells dominant negative *P53* mutations have been identified that confer a selective advantage to the cells in culture ^14^. Based on our *in vitro* mutation accumulation results we predict that these mutations occur once in every ∼5.7*10^8^ iPS cells (figure 3a). As another example, we focused on the *BRAF*^V600E^ oncogenic mutation, which is found in ∼10% of colorectal cancers^29^. In intestinal ASCs this mutation occurs once in every 1.1 *10^10^ stem cells (figure 3a). The chance for having a *BRAF*^V600E^ mutation in the total cell population is 0.018, when 10^8^ intestinal ASCs are produced *in vitro*. This chance is equivalent to the probability that a *BRAF*^V600E^ mutation occurs in any colon stem cell in 112 days of adult age (figure 3b).

**Figure 3:**
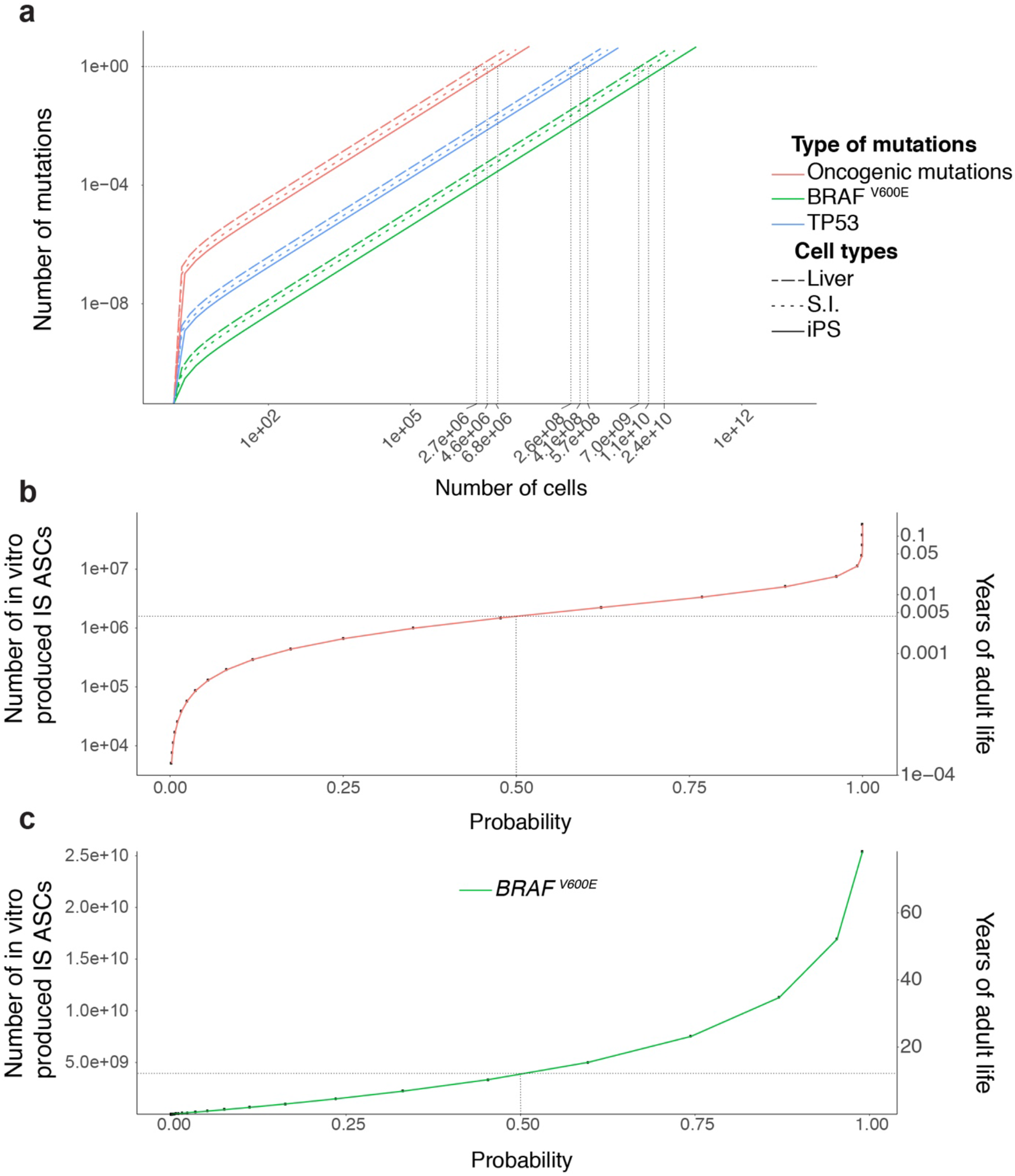
Modelling the oncogenic mutations in *in vitro* cultured stem cells. **a**, The number of oncogenic mutations as a function of the number of in vitro produced cells. **b**, The probability of oncogenic mutations (x-axis) as a function of the number of in vitro produced ASCs (primary y-axis) and as a function of years of adult life (secondary y-axis). **c**, The probability of *BRAF*^V600E^ mutations (x-axis) as a function of the number of in vitro produced ASCs (primary y-axis) and as a function of years of adult life (secondary y-axis).

## Discussion

Many studies have described genetic abnormalities in stem cells during *in vitro* culture and/or derivation. These findings seem to contrast with the observation that stem cells have a higher activity of DNA repair pathways and repair damaged DNA more efficiently than differentiated cell types ^30–32^. To systematically investigate genome stability of stem cells, and quantify mutation accumulation, we have applied whole genome sequencing of iPS cells and ASCs to identify genetic aberrations that were acquired during the culturing period in between two clonal steps.

An important distinction from previous studies is that this approach enables to discriminate mutations that originate in culture from those that have an *in vivo* origin or arise as a result of the derivation process. Previously, we discovered that, *in vivo*, ASCs acquire ∼40 mutations per year ^18^, so for specific experimental setups the observed mutations in cultured cells does depend on the age of the donor. By our experimental approach we excluded these confounding factors and demonstrate that the *in vitro* mutation rates for ASCs are ∼40-fold higher than *in vivo*. For tumor-derived stem cells ^33^, mutation rates could be even higher due to pre-existing increased mutation rates in tumor cells. However, the *in vivo* number should be considered in a life-long context (decades) over a large population of stem cells, while the *in vitro* number relates to the duration of culturing (weeks or months) and the total number of cells required.

Our mutational signature analyses suggest that iPS cells, liver stem cells, and intestinal stem cells experience distinct intrinsic mutational processes. For example, the intestinal stem cells showed a clear transcriptional strand-bias for T to A mutations that was not observed in the other stem cell types. Likewise, signature 5 contributed to all iPS cell subclones. In agreement with signature 5, the iPS cells showed strong transcriptional strand bias for T to C substitutions. These findings suggest differential activity of DNA repair pathways between stem cell types. Furthermore, our results show that iPS cells acquire fewer SNVs, indels, and SVs per population doubling than ASCs. This suggests that iPS cells experience lower levels of DNA damage or have higher activity of DNA repair pathways multiple DNA repair pathways. Previous studies have indeed demonstrated that human pluripotent stem cells exhibit high activity of DNA repair pathways leading to enhanced capacity to repair damaged DNA ^32,34^. However, considering the stem-cell specific mutation rates, mutation spectra, and genomic distribution, iPS cells acquire only slightly fewer oncogenic mutations than ASCs. However, the depletion of mutations from promoter regions is stronger in the ASCs than in the iPS cells. Thus, human iPS cells are more protected from harmful coding sequence changes but more vulnerable to detrimental mutations in regulatory promoter regions.

There is a strong correlation between the number of mutations and the number of cells produced. As a consequence, the majority of the mutations arise during the last population doubling and are therefore present at very low frequencies, but also remain undetected in pool-based approaches. It is therefore impossible to rule out the risk that a population of cells that are being transplanted is devoid of any oncogenic mutation. This obviates the need to rely on accurate risk assessment for oncogenic mutations to occur. Our examples illustrate that oncogenic mutations will inevitably arise during culture as they do *in vivo*, but the risk for a specific oncogenic mutation, such as *BRAF*^V600E^, is low. We therefore think that the observed *in vitro* mutation rates should not impede their future use in regenerative medicine. However, to avoid unnecessary mutation accumulation we recommend to minimize the time in culture.

The *in vitro* mutation spectra of all three stem cell types are characterized by high numbers of C to A transversions, which are probably caused by ROS. High ROS-levels can cause oxidative damage to guanine, resulting in the formation of 8-oxoguanine. Incorporation of dAMP opposite 8-oxoguanine escapes exonucleolytic proofreading by DNA polymerases, generating C to A transversions during the next round of replication ^27^. Lowering the levels of ROS in culture may thus be a suitable intervention to reduce the number of C to A changes and the overall mutation rates, for all three stem cell types. The levels of DNA damage are indeed reduced when human pluripotent stem cells are cultured under low oxygen tension, but the effect on mutation spectrum remains to be experimentally validated ^3^.

Taken together, the experimental approach of the current study provides a suitable framework for studies on the effects of mutagenic environmental factors during in vitro culture and allows for further optimization of culture conditions to eventually reflect *in vivo* mutation rates and to establish safe *in human* applications of these powerful cellular tools.

## Material and Methods

### Stem cell culture

All tissue culture products were from ThermoFisher scientific, unless stated otherwise. Integration-free human iPS cells were established from urinary cells using Sendai virus. Collection and culture of primary urinary cells was performed as previously described ^35^. In short, urine was collected from a healthy male volunteer and divided over 50ml tubes followed by centrifugation. The cell pellets were washed with PBS supplemented with antibiotics. The cell pellets were resuspended in primary medium containing DMEM/high glucose and Ham’s F12 nutrient mix, supplemented with 10% (vol/vol) FBS, pen/strep, Renal Cell Growth Medium SingleQuots™ (Lonza) 2.5 μg ml−1 amphotericin B. The cells were plated onto gelatin-coated plates and incubated at 37°C in a humidified atmosphere and 5% CO_2_. Medium was refreshed daily. 96 hours after plating the medium was switched to Renal Cell Growth Medium supplemented with the SingleQuots™. At near confluency the cells were split to new gelatin coated plates using TrypLE Express. Urinary cells were subsequently reprogrammed by transduction with Sendai virus carrying human Klf4, Oct3/4, Sox2, and C-Myc using the CytoTune™ iPS 2.0 Sendai Reprogramming Kit according to the manufacturer’s protocol. Approximately 10 days after transduction, individual iPS colonies were manually picked and cultured in E8 medium on Geltrex-coated plates ^36^. RNA-seq analysis confirmed that the iPS cells closely resembled human embryonic stem cells (supplemental figure 6). Clonal steps were performed by limiting dilution of a single cell suspension in Geltrex-coated 96-well plates. To enhance cell survival after the clonal steps of the iPS cells, E8 medium was supplemented with RevitaCell™. In between two clonal steps, the iPS cells were cultured for 3 months to accumulate mutations. After a three-month culture, a second clonal step was performed. The resulting clones were expanded until enough material was produced for whole genome sequencing. To filter out germline variants, we used a cell pellet from the pre-clonal bulk culture as reference sample.

For ASCs, we used previously established human liver and intestinal stem cell lines that were cultured under previously described conditions ^18,26^. In short, liver stem cell lines were cultured in Advanced DMEM/F12 with HEPES, Glutamax, and Penicillin Streptomycin. This medium was further supplemented with N2, B27, 1.25 mM N-Acetylcysteine (Sigma), 10 nM gastrin (Tocris) Primocin (Invivogen), and the following growth factors: 50 ng/ml EGF (Peprotech), 10% RSPO1 conditioned media (homemade), 100 ng/ml FGF10 (Peprotech), 25 ng/ml HGF (Peprotech), 10 mM Nicotinamide (Sigma), 5 uM A83.01 (Tocris), and 10 μM FSK (Tocris). For the first 3 days after each clonal step the medium contained 25 ng/ml Noggin (Peprotech), 30% Wnt3a conditioned medium(homemade made 37), 10 μM Y27632 (Abmole) and stem cell cloning recovery solution (Stemgent). Intestinal stem cell lines were cultured in Advanced DMEM/F12 with HEPES, Glutamax, and Penicillin Streptomycin. This medium was further supplemented with 50% Wnt3a conditioned medium, 20% R-Spondin conditioned medium, B27, 1.25 mM N-Acetylcysteine, 10 mM Nicotinamide, Primocin, 0.5mM A83–01 (Tocris), 30mM SB202190 (Sigma), recombinant human Noggin (Peprotech) and 50ng/ml recombinant human EGF (Peprotech). For the first 3 days after each clonal step the intestinal organoid medium was supplemented with 10 μM Y27632

To make clonal stem cell lines, organoids were enzymatically dissociated and the resulting single cell suspension was FACS-sorted to remove all doublets. The single cell suspension was resuspended in Matrigel (Corning) or BME (Pathclear) for intestinal stem cells and liver stem cells respectively and plated as a limiting dilution series. After 2–3 weeks, individual organoids were manually picked and mechanically fragmented and plated under regular culture conditions. Clonal lines were further cultured for another 53–146 days after which a second clonal step was performed. The resulting subclones were expanded until enough material was available for whole genome sequencing. Per stem cell line, a blood sample of the same also donor was also sequenced to enable filtering for germline variants.

### DNA/RNA isolation, library prep and sequencing

DNA was isolated manually using the genomic tip 20-G kit (Qiagen) or automated using the Qiasymphony (Qiagen). From 200 ng genomic DNA, DNA libraries were generated for Illumina sequencing using standard protocols (Illumina). Libraries were sequenced 2 × 100 bp paired-end to 30X base coverage with the Illumina HiSeq Xten at the Hartwig Medical Foundation. For RNA-sequencing, human iPS cells, ES cells, liver ASCs and Intestinal ASCs were collected in Trizol and subjected to total RNA isolation with the QiaSymphony SP using the QiaSymphony RNA kit (Qiagen, 931636). Subsequently, 50 ng total RNA was used to prepare mRNA sequencing libraries using the Illumina Neoprep TruSeq stranded mRNA library prep kit (Illumina, NP-202–1001), followed by paired-end sequencing (2 × 75 bp) of the RNA libraries on a Nextseq500 to > 20 million reads per sample. All sequencing data has been deposited at the European Genome-phenome Archive (http://www.ebi.ac.uk/ega/) under accession numbers EGAS00001002955, EGAS00001000881 and EGAS00001001682.

### Sequencing analysis

RNA sequencing reads were mapped to the human reference genome GRCh37 with STAR v.2.4.2a ^38^ and the BAM-files were indexed using Sambamba v0.5.8. Reads were counted using HTSeq-count 0.6.1p1 and read counts were normalized using DESeq v1.28.0. Non-supervised hierarchical clustering was performed using DESeq2.0 v1.20. For DNA sequencing read mapping and data-preprocessing was performed as previously described ^16^.

We determined the ‘callable regions’ of the genome for each sample that was sequenced based on coverage and read quality for each genomic position. Variants located in regions that were not callable in the reference sample were excluded from the analysis. For each sample, all mutations were normalized to the callable genome to determine the total number of mutations per whole genome (supplementary table 1). To identify all SNVs of the clonal cultures we used our data analysis pipeline that enables the identification of somatic SNVs with a confirmation rate of ∼91%^18^. In short, we performed basic protocols 1–2 of the Genome Analysis Toolkit (GATK) best practices workflow for germline single-nucleotide polymorphisms (SNPs) and indels in whole genomes to identify SNVs^39^. We subsequently generated a catalogue of high-quality *in vitro* induced SNVs using a custom Single Nucleotide Variant Filtering pipeline (SNVFI available at https://github.com/UMCUGenetics/SNVFI). The SNV call set is further filtered on the basis of several quality parameters and dbSNP v137.b37 ^40^. In addition, positions that were found to be variable in at least three unrelated individuals were excluded as these represent either unknown SNPs or recurring sequencing and/or calling artefacts ^16^. Furthermore, events with any evidence in the reference sample (alternative depth >0) were excluded. Clonality of the cultures was verified by a distribution around 0.5 of the variant allele frequencies (VAFs) of the mutations. By filtering against all mutations with allele frequencies below 0.3 we excluded all mutations that arose during the culture period after the clonal steps. To identify all the mutations that originated in the culture period in between both clonal steps, the mutations identified in the second clonal culture were filtered for those present in the bulk and the first clonal culture. The resulting number of SNVs was divided by the population doublings, to obtain the point mutation rate for each stem cell type. Welch Two Sample t-tests were performed to determine whether the mutation rates differ significantly between the stem cell types. To assess the presence of the mutations within genes and to predict their effect, the SNVs were annotated using SnpEff ^41^. SNVs mutational profiles and genomic distributions were obtained using the R package MutationalPatterns ^18^. The COSMIC mutational signatures were downloaded from COSMIC website (https://cancer.sanger.ac.uk/cosmic) and their relative contribution to the total number of mutations was calculated for each sample. Genomic and transcriptional features were extracted from the hg19 assembly downloaded from the UCSC Table Browser (https://genome.ucsc.edu/cgi-bin/hgTables), while epigenetic status datasets were downloaded from the ENCODE website (https://cancer.sanger.ac.uk/cosmic) and their relative contribution to the total number of mutations was calculated for each sample. Genomic and transcriptional features were extracted from the hg19 assembly downloaded from the UCSC Table Browser (https://genome.ucsc.edu/cgi-bin/hgTables), while epigenetic status datasets were downloaded from the ENCODE website (https://www.encodeproject.org/) as BED files; then the SNVs distribution and enrichment or depletion within these regions was calculated.

Unfiltered indel catalogs were extracted from the germline GATK calls acquired in ‘Single nucleotide variants’. We only considered indels within the callable autosomal genome of the clone, subclone, and control sample. We subsequently excluded indels that overlap with a SNV, indels that have a dbSNP ID and no COSMIC ID, and indels that are present on a blacklist of 3 unrelated samples (BED file available upon request). Furthermore, we only considered indels with a GATK quality score of at least 250, with a filter ‘PASS’ from VariantFiltration, with a coverage of at least 20x in clone, subclone, and control sample, and with a sample-specific genotype quality of at least 99 in clone and subclone. We subsequently excluded indels with any evidence of an alternative call in the control sample and/or any of the other samples, with a VAF > 0.3 in the clone, and with a VAF < 0.3 in the subclone. The Indels were inspected manually in IGV, to excluded false positives.

We extrapolated the number of indels to the human autosomal genome. Subsequently the number of indels was divided by the number of population doublings, to obtain the indel mutation rate for each stem cell type. An ANOVA was performed to determine whether indel mutation rates differ significantly between stem cell types. To test for enrichment and depletion of indels within genes, we used one-sided binomial test with MutationalPatterns ^42^. Finally, we used SnpEff variant annotation to predict the effect of the mutations on the genes ^41^.

Copy-number variants (CNVs) were called using Control-FREEC^43^ and filtered on the basis of size and evidence in the other samples. For calling structural variation we used manta v1.1.0 with default settings, and annotated the calls using break-point-inspector version [1] from commit 8d30505dfab219e367a6e5d7d3f2e6ec74877e75. We manually inspected all variants containing the “PASS” filter option using IGV to determine true-positive variant calls, and whether the structural variant accumulated in vitro.

### Modelling of *in vitro* mutation accumulation

To calculate population doubling time we used the split ratios to calculate the number of cells we would have obtained under maximum expansion after a culture period of 44 days. Next, we applied the formula 
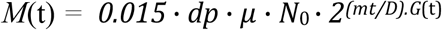
where T is the incubation time in any units, Xb is the cell number at the beginning of the incubation time and Xe is the cell number at the end of the incubation time. To calculate the number of mutations in the protein coding fraction of the genome we applied the formula: 
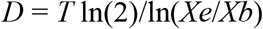
where *M*(t) is the number of mutations at timepoint t, 0.015 is the coding fraction of the whole genome and *dp* is the degree of depletion in the CDS, *μ* is the number of mutations per cell cycle, *N_(0)_* is the number of initial cells, *mt* is the cell cycle length, and *G(t)* is the number of generations at timepoint t. For ASCs we used a cell cycle length of ∼26h as determined in intestinal stem cells (data not shown) and for iPS cells we applied a cell cycle length of ∼18h^44^.

We used a list of oncogenic mutations in driver genes from Tamborero et al.^28^ to calculate the number of mutations in the coding sequence activate cancer driver genes. The probabilities *P* that mutation types (*C* → *A, C* → *G, C* → *T, T* → *A, T* → *C* and *T* → *G*) happen in the genome can be derived from the mutation spectrum of the cell type. To calculate the number of oncogenic mutations as a function of the number of cells, we applied the following formula: 
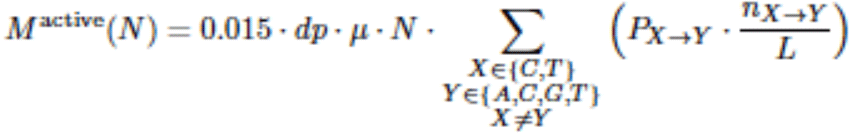
where M^active^ is number of mutations that activate driver genes, dp is depletion in CDS, *μ* is the mutation rate, N is number of cells, P_X>Y_ is chance on X>Y mutation based on the mutation spectrum, n_X>Y_ is the number of positions where X>Y mutation result in oncogene activation and L is the length of CDS.

To calculate the probability for an oncogenic mutation as a function of the number of cells we applied the following formula, where Z is number of activating mutations. 
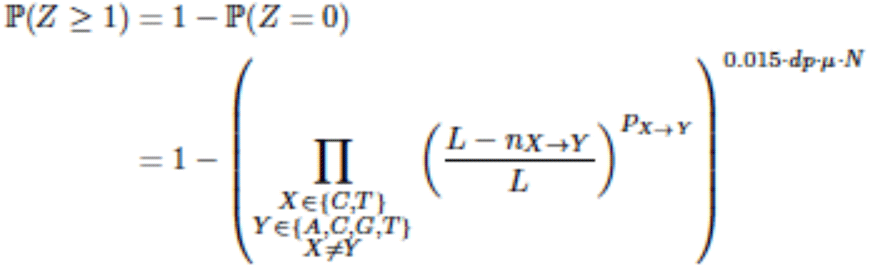

To calculate the probability for an oncogenic mutation *in vivo* in 10^8^ colon stem cells as a function of the number of years (t) in adult life we applied the following formula: 
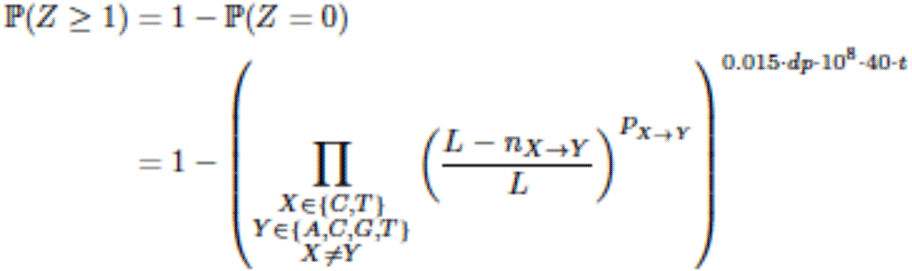

## Acknowledgments

The authors thank the USEQ for sequencing support and the UBEC for bioinformatical support. This work was financially supported by the NWO Gravitation Program Cancer Genomics.nl and the NWO/ZonMW Zenith project 93512003 to E.C.

## Author contributions

M.J. E.C. and E.K. wrote the manuscript. N.B., R.v.B., M.J., J.K., E.K. performed wet-lab experiments. M.J., R.J., A.V.H., M.L. and E.K. performed bioinformatical analyses. R.J., A.V.H., B. vd R., and E.K performed the mathematical modelling. S.B. provided bioinformatical support. R.v.B., E.C., M.J., and E.K. were involved in the conceptual design of the study. E.C. supervised the study.

## Supplementary figures

**Supplementary figure 1:**
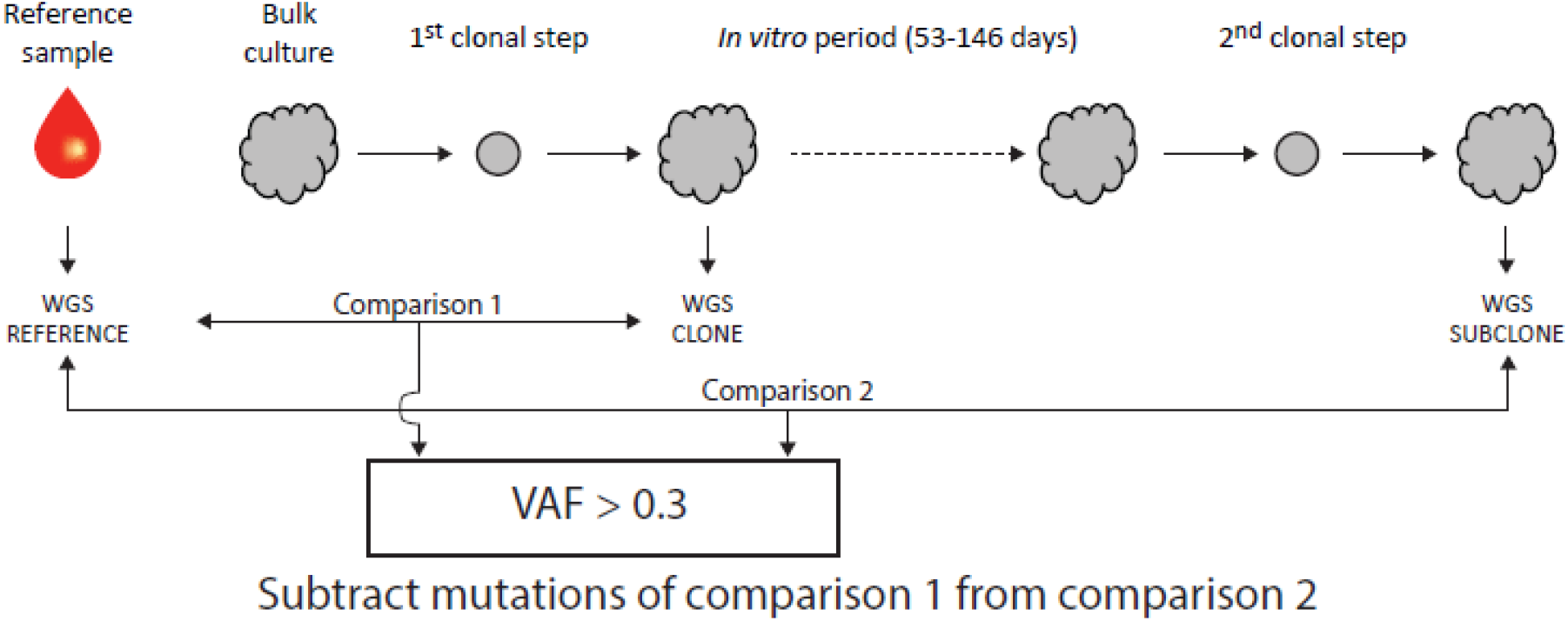
Experimental set-up. Schematic overview of the experimental setup to determine *in vitro* accumulated mutations in individual human iPS cells, liver stem cells, and intestinal stem cells. Clonal stem cell lines were cultured for ∼2–5 months during which mutations were allowed to accumulate. At the end of the culture period a 2^nd^ clonal step was performed and the derivative subclones were expanded until enough DNA could be isolated for WGS analysis. A biopsy or bulk culture was used as a reference sample to determine and exclude all germline variants.

**Supplementary figure 2:**
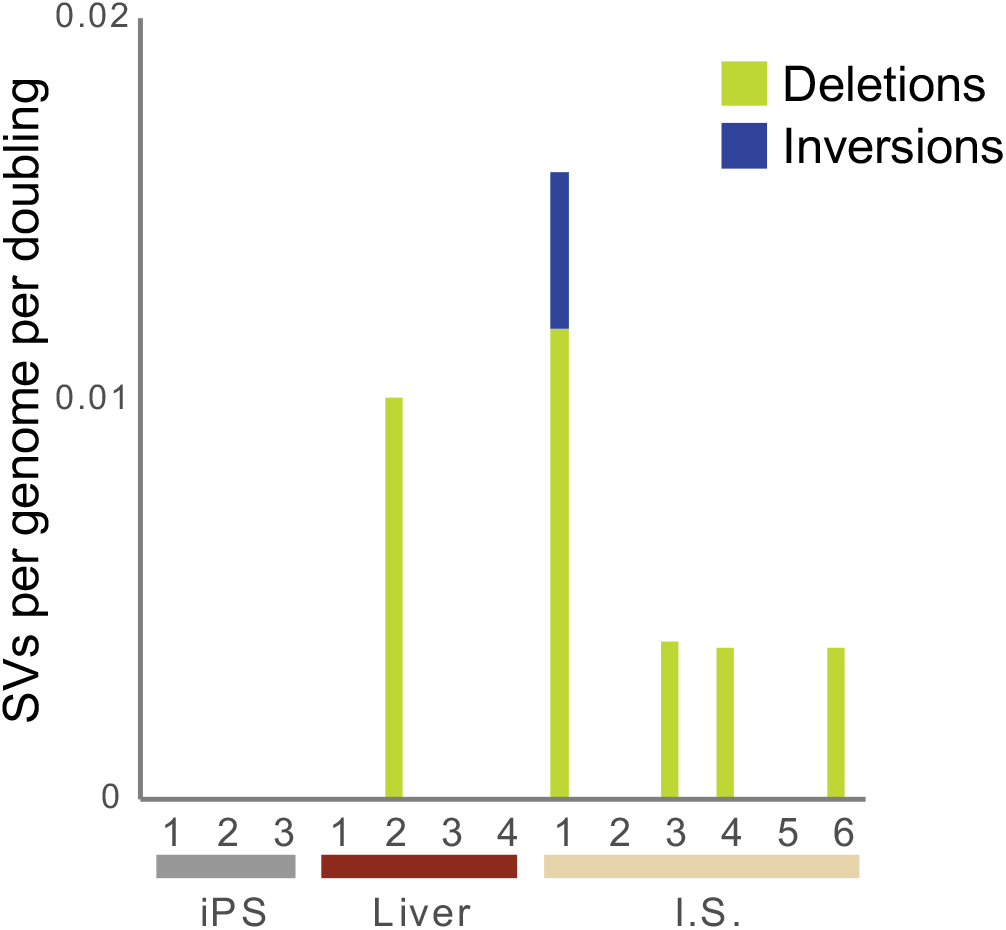
*In vitro* accumulation of structural variation in iPS cells, liver stem cells, and intestinal stem cells.

**supplemental figure 3.**
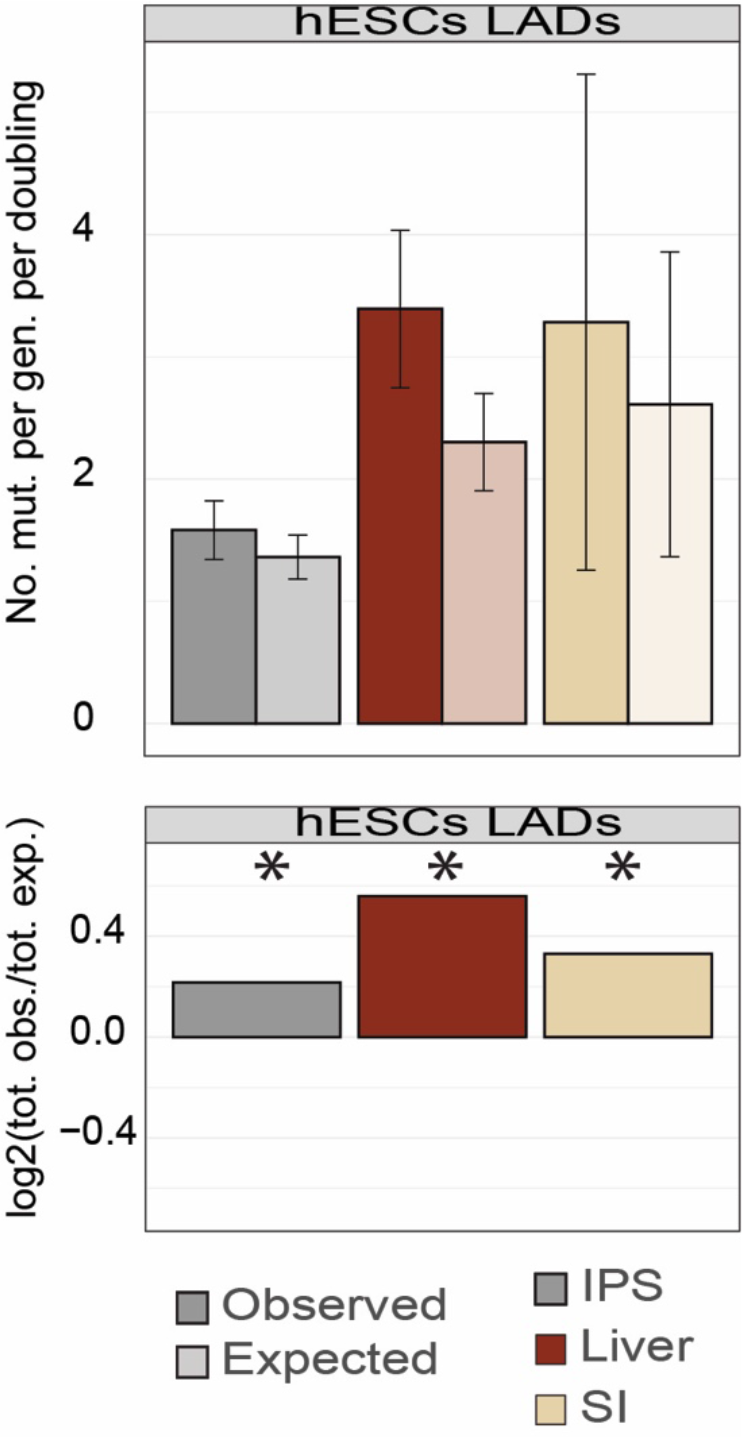
Genomic distribution. Observed versus expected amount of mutations at laminin associated domains. Asterisks denote significant differences between observed and expected (binomial test).

**supplemental figure 4:**
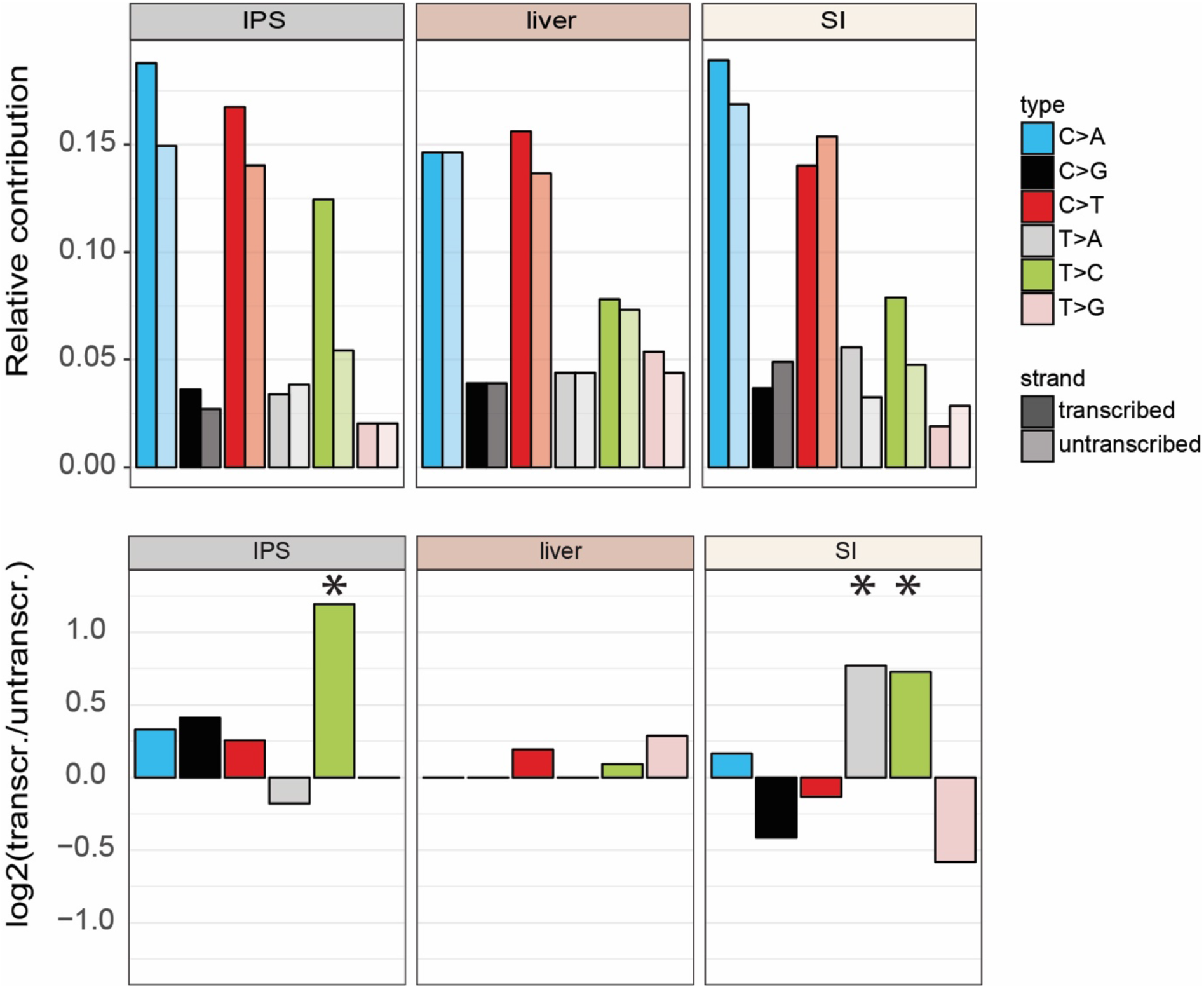
Transcriptional strand bias of *in vitro* accumulated mutations. Asterisks denote significant differences between observed and expected (binomial test).

**supplemental figure 5:**
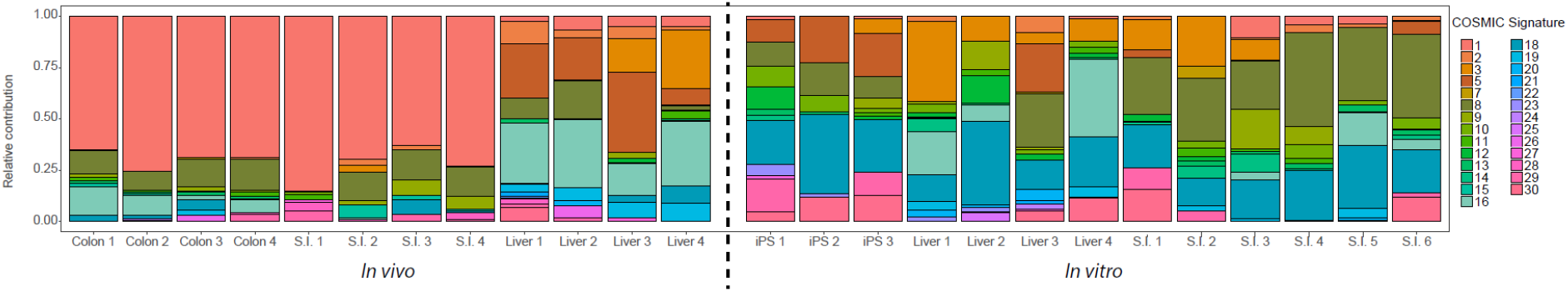
Relative contribution of the COSMIC signatures to the *in vivo* and *in vitro* mutation accumulation in stem cells.

**supplemental figure 6:**
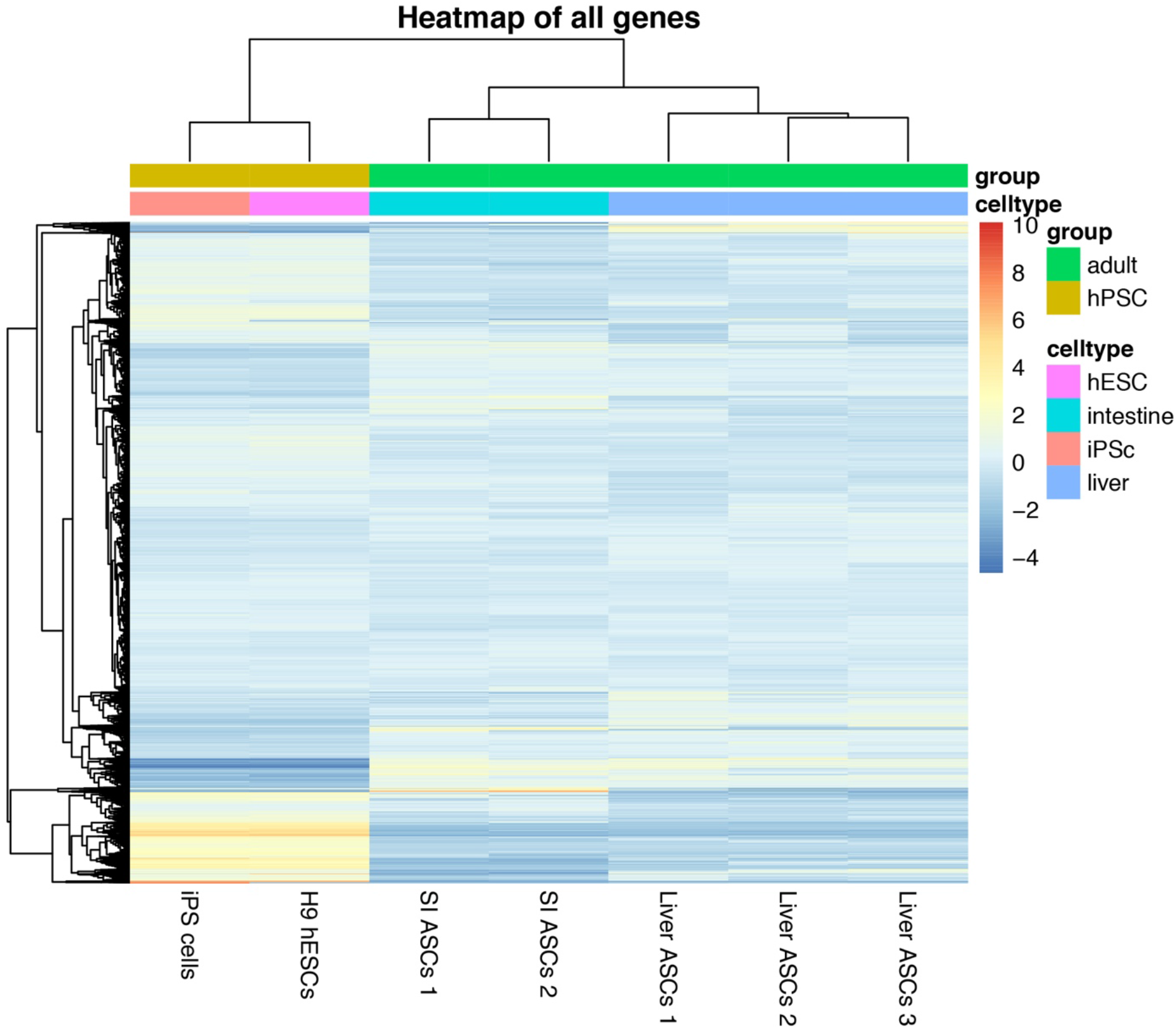
Heatmap and unsupervised hierarchical clustering of RNA-seq data for all genes for human iPS cells, human ES cells, human liver stem cells and human intestinal stem cells.

## Supplementary tables

**supplementary table 1:** Days in culture, callable genome, number of mutations, and the number of mutations corrected for the callable genome for each subclone.

| Cell type | Subclone | Days in culture | Callable | Total genome size | Callable fraction | Uncorrected indels | Corrected for callable genome | Indels/day | Indels/month | Doubling time (h) | Indels/doubling |
| --- | --- | --- | --- | --- | --- | --- | --- | --- | --- | --- | --- |
| Liver organoids | 1 | 53 | 1915892864 | 2881033286 | 0.67 | 8 | 12.03 | 0.23 | 6.81 | 45.82 | 0.43 |
| Liver organoids | 2 | 53 | 1633429010 | 2881033286 | 0.57 | 7 | 12.35 | 0.23 | 6.99 | 45.82 | 0.44 |
| Liver organoids | 3 | 92 | 1613568163 | 2881033286 | 0.56 | 5 | 8.93 | 0.10 | 2.91 | 45.82 | 0.19 |
| Liver organoids | 4 | 88 | 1765097010 | 2881033286 | 0.61 | 11 | 17.94 | 0.20 | 6.12 | 45.82 | 0.39 |
| iPS cells | 1 | 92 | 2560647952 | 2881033286 | 0.89 | 14 | 15.75 | 0.17 | 5.14 | 23.08 | 0.16 |
| iPS cells | 2 | 92 | 2568349174 | 2881033286 | 0.89 | 15 | 16.84 | 0.18 | 5.49 | 23.08 | 0.18 |
| iPS cells | 3 | 92 | 2551<br>9309<br>84 | 288103<br>3286 | 0.89 | 9 | 10.16 | 0.11 | 3.31 | 23.08 | 0.11 |
| Intestinal organoids | 1 | 140 | 2106<br>5701<br>46 | 288103<br>3286 | 0.73 | 14 | 19.15 | 0.14 | 4.10 | 44.38 | 0.25 |
| Intestinal organoids | 2 | 140 | 2086<br>1743<br>48 | 288103<br>3286 | 0.72 | 34 | 46.96 | 0.34 | 10.06 | 44.38 | 0.62 |
| Intestinal organoids | 3 | 140 | 1992<br>1904<br>47 | 288103<br>3286 | 0.69 | 28 | 40.52 | 0.29 | 8.68 | 44.38 | 0.54 |
| Intestinal organoids | 4 | 146 | 1840<br>3247<br>04 | 288103<br>3286 | 0.64 | 16 | 25.04 | 0.17 | 5.15 | 44.38 | 0.32 |
| Intestinal organoids | 5 | 146 | 2016<br>9469<br>17 | 288103<br>3286 | 0.70 | 23 | 32.86 | 0.23 | 6.75 | 44.38 | 0.42 |
| Intestinal organoids | 6 | 146 | 1804<br>3793<br>89 | 288103<br>3286 | 0.63 | 17 | 27.16 | 0.19 | 5.58 | 44.38 | 0.34 |

**supplementary table 2:** All structural variants and their genomic locations.

| Tissue | Subclone | SVTYPE | ORIENTATION | MANTABP1 | MANTABP2 | SV size | Affected genes | Intronic/exonic | Common fragile site |
| --- | --- | --- | --- | --- | --- | --- | --- | --- | --- |
| Liver | 2 | DEL | INNIE | 14:88744986 | 14:88745125 | 139 | KCNK10 | Intronic | NA |
| S.I. | 1 | DEL | INNIE | 3:60406136 | 3:60433613 | 27477 | FHIT | Intronic | CFS 3B |
| S.I. | 1 | DEL | INNIE | 3:60434816 | 3:60531880 | 97064 | FHIT | Intronic | CFS 3B |
| S.I. | 1 | DEL | INNIE | 3:60612721 | 3:60612773 | 52 | FHIT | Intronic | CFS 3B |
| S.I. | 1 | INV | TANDEM_LEFT | 6:75110810 | 6:75114894 | 4084 | NA | NA | NA |
| S.I. | 3 | DEL | INNIE | 16:78997362 | 16:79066865 | 69503 | WWOX | Intronic | CFS 16D |
| S.I. | 4 | DEL | INNIE | 1:7136482 | 1:7136533 | 51 | CAMTA1 | Intronic | NA |
| S.I. | 6 | DEL | INNIE | X:121224087 | X:121225190 | 1103 | NA | NA | NA |

**supplementary table 3:**
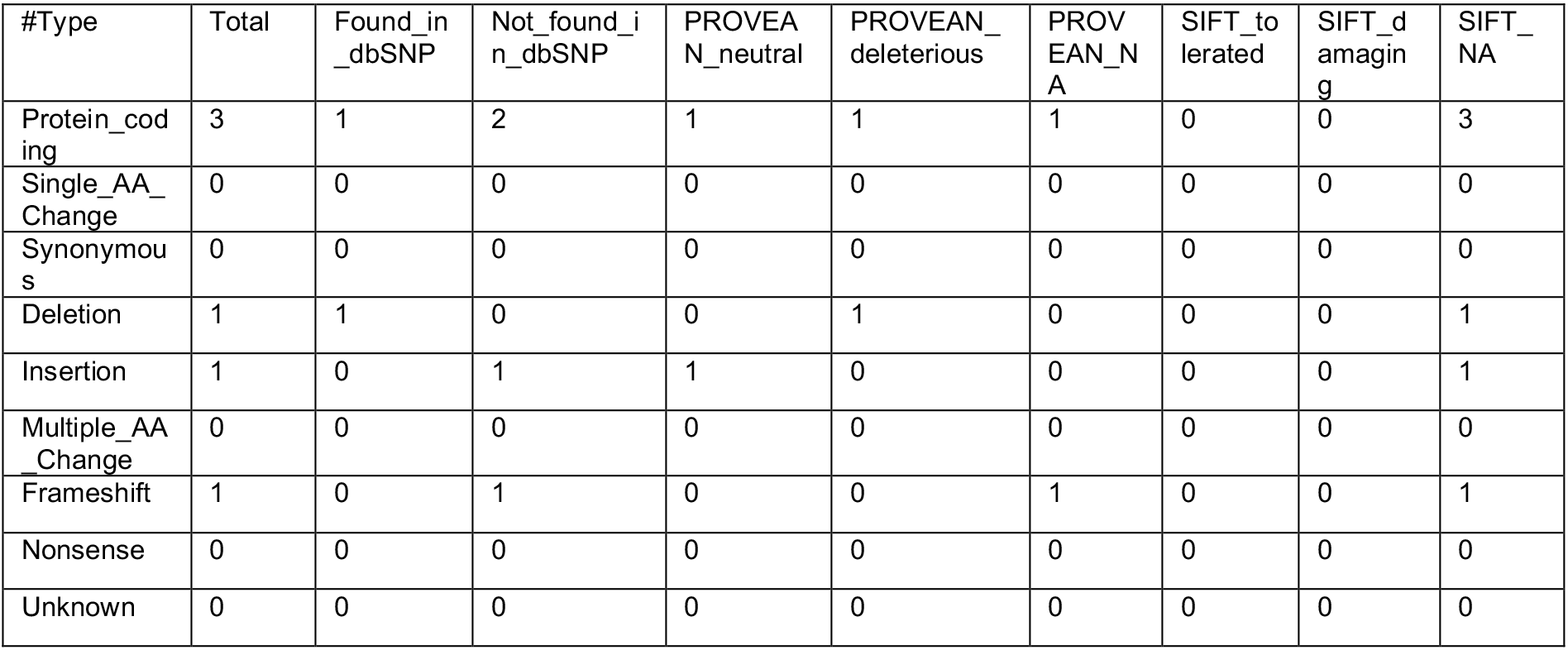
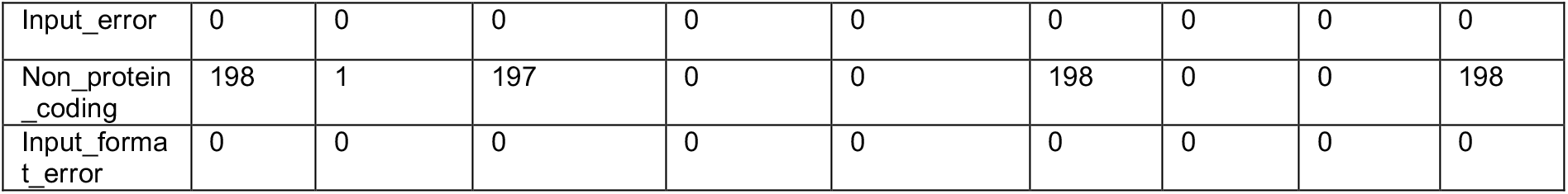
PROVEAN analysis results for all indels.

**supplementary table 4:** all mutations in the coding part of the genome.

| Gene | mut type | Subclone | stem cell type |
| --- | --- | --- | --- |
| ZMYM1 | missense | 1 | iPS |
| LAMC3 | missense | 1 | iPS |
| GYS2 | missense | 2 | iPS |
| SLC39A6 | missense | 2 | iPS |
| LDLR | missense | 2 | iPS |
| RSL24D1 | missense | 3 | liver |
| CEP135 | missense | 3 | liver |
| XDH | missense | 1 | small intestine |
| CYP11B1 | missense | 1 | small intestine |
| WDR59 | missense | 6 | small intestine |
| KIF2B | missense | 6 | small intestine |
| SPACA1 | missense | 4 | small intestine |
| LHFPL3 | missense | 4 | small intestine |
| DISP2 | missense | 6 | small intestine |
| MYH14 | synonymous | A3 | iPS |
| CRB1 | synonymous | B3 | iPS |
| BAI2 | synonymous | 1 | liver |
| SYNE1 | synonymous | 3 | liver |
| OR2G3 | synonymous | 1 | small intestine |
| SPTBN4 | synonymous | 2 | small intestine |
| FSIP2 | synonymous | 2 | small intestine |
| DNAH1 | synonymous | 2 | small intestine |
| RHOBTB1 | synonymous | 5 | small intestine |
| PKHD1L1 | synonymous | 6 | small intestine |

